# Early metabolic features of genetic liability to type 2 diabetes: cohort study with repeated metabolomics across early life

**DOI:** 10.1101/767756

**Authors:** Joshua A. Bell, Caroline J. Bull, Marc J. Gunter, David Carslake, George Davey Smith, Nicholas J. Timpson, Emma E. Vincent

## Abstract

**Background:** Type 2 diabetes develops for many years before diagnosis. We aimed to reveal early metabolic features characterising liability to adult disease by examining genetic liability to adult type 2 diabetes in relation to detailed metabolic traits across early life.

**Methods and Findings:** Data were from up to 4,761 offspring from the Avon Longitudinal Study of Parents and Children cohort. Linear models were used to examine effects of a genetic risk score (GRS, including 162 variants) for adult type 2 diabetes on 4 repeated measures of 229 traits from targeted nuclear magnetic resonance (NMR) metabolomics. These traits included lipoprotein subclass-specific cholesterol and triglyceride content, amino and fatty acids, inflammatory glycoprotein acetyls, and others, and were measured in childhood (age 8y), adolescence (age 16y), young-adulthood (age 18y), and adulthood (age 25y). For replication, two-sample Mendelian randomization (MR) was conducted using summary data from genome-wide association studies of metabolic traits from NMR in an independent sample of adults (N range 13,476 to 24,925; mean (SD) age range 23.9y (2.1y) to 61.3y (2.9y)). Among ALSPAC participants (49.7% male), the prevalence of type 2 diabetes was very low across time points (< 5 cases when first assessed at age 16y; 7 cases (0.4%) when assessed at age 25y). At age 8y, type 2 diabetes liability (per SD-higher GRS) was associated with lower lipids in high-density lipoprotein (HDL) particle subtypes – e.g. −0.03 SD (95% CI = −0.06, −0.003; P = 0.03) for total lipids in very-large HDL. At age 16y, associations remained strongest with lower lipids in HDL and became stronger with pre-glycemic traits including citrate (−0.06 SD, 95% CI = −0.09, −0.02; P = 1.41×10^−03^) and with glycoprotein acetyls (0.05 SD, 95% CI = 0.01, 0.08; P = 0.01). At age 18y, associations were stronger with branched chain amino acids including valine (0.06 SD; 95% CI = 0.02, 0.09; P = 1.24×10^−03^), while at age 25y, associations had strengthened with VLDL lipids and remained consistent with previously altered traits including HDL lipids. Results of two-sample MR in an independent sample of adults indicated persistent patterns of effect of type 2 diabetes liability, with higher type 2 diabetes liability positively associated with VLDL lipids and branched chain amino acid levels, and inversely associated with HDL lipids – again for large and very large HDL particularly (−0.004 SD (95% CI = −0.007, −0.002; P = 8.45×10^−04^) per 1 log odds of type 2 diabetes for total lipids in large HDL). Study limitations include modest sample sizes for ALSPAC analyses and limited coverage of protein and hormonal traits; insulin was absent as it is not quantified by NMR and not consistently available at each time point. Analyses were restricted to white-Europeans which reduced confounding by population structure but limited inference to other ethnic groups.

**Conclusions:** Our results support perturbed HDL lipid metabolism as one of the earliest features of type 2 diabetes liability which precedes higher branched chain amino acid and inflammatory glycoprotein acetyl levels. This feature is apparent in childhood as early as age 8y, decades before the clinical onset of disease.

**Author summary:** *Why was this study done?:*

- Type 2 diabetes develops for many years before diagnosis. Clinical disease is characterised by numerous metabolic perturbations that are detectable in circulation, but which of these reflect the developmental stages of type 2 diabetes – as opposed to independent causes of type 2 diabetes or markers of other disease processes – is unknown. Revealing traits specific to type 2 diabetes development could inform the targeting of key pathways to prevent the clinical onset of disease and its complications.
- Genetic liability to type 2 diabetes is less prone to confounding than measured type 2 diabetes or blood glucose and may help reveal early perturbations in the blood that arise in response to type 2 diabetes liability itself.

*What did the researchers do and find?:*

- We examined effects of genetic liability to adult type 2 diabetes, based on a genetic risk score including 162 variants, on detailed metabolic traits measured on the same individuals across four stages of early life – childhood (age 8y), adolescence (age 16y), young-adulthood (age 18y), and adulthood (age 25y).
- We found that higher type 2 diabetes liability was associated most consistently across ages with lower lipid content in certain subtypes of HDL particles. Effects were more gradual on higher lipid content in VLDL particles and on higher branched chain amino acid and inflammatory glycoprotein acetyl levels.

*What do these findings mean?:*

- Signs of type 2 diabetes liability are detectable in the blood in childhood, decades before the disease becomes noticeable. These signs, taken to reflect the early features of, or coincident with, disease, likely involve lower lipid content in HDL particles, followed by higher levels of branched chain amino acids and inflammation.
- Genetic risk scores for adult diseases can be integrated with metabolic measurements taken earlier in life to help to reveal the timing at which signs of disease liability become visible and the traits most central to its development.

## Introduction

Type 2 diabetes is a metabolic disease affecting more than 400 million people globally [1]. Its incidence is driven largely by increased adiposity [2], a strong causal risk factor [3,4], yet the difficulty of achieving and maintaining non-surgical weight loss makes disease management a lifelong and expensive task [5]. This is particularly problematic considering that potentially half of those living with type 2 diabetes are undiagnosed and the future burden is expected to be greatest in lower income countries [1]. There is therefore a clear need to minimise the impact of type 2 diabetes on individuals and populations, and this requires biological understanding of the disease at its very earliest stages.

Type 2 diabetes is typically diagnosed when blood glucose levels exceed 7 mmol/l in the fasting state or 11.1 mmol/l in the post-challenge state, or when glycated haemoglobin levels exceed 6.5% [6], yet glucose spikes relatively late in the disease process. Repeat clinical measures from the Whitehall II cohort study suggest that insulin sensitivity starts declining a decade before glucose changes are detectable [7]. Cohort studies with targeted metabolomic measurements also observe associations of numerous subclinical traits with lower insulin sensitivity including higher branched chain and aromatic amino acid concentrations; higher fatty acid and inflammatory glycoprotein concentrations; and elevated lactate and pyruvate levels [8-11]. Relations with ketone bodies are less clear, since higher levels are associated with both higher insulin sensitivity [9] and higher type 2 diabetes risk [12]. Hyperglycemia also associates strongly with cholesterol and triglyceride content across lipoprotein subclasses [13,14]. Whether such trait alterations reflect true developmental stages of type 2 diabetes is unclear because of inherent confounding by other disease processes.

An alternative approach is to examine genetic liability to type 2 diabetes – rather than measured type 2 diabetes or hyperglycaemia – in relation to metabolic traits, which could help identify perturbations specific to the development of type 2 diabetes itself – i.e. its early metabolic features [15,16]. The few studies that have investigated this suggest effects of type 2 diabetes liability on cholesterol and triglycerides in high-density lipoprotein (HDL) and very-low density lipoprotein (VLDL) particles [17], amino acids [18,19], and ketone bodies [12]. Most used a small set of genetic variants from early genome-wide association studies (GWAS) and did not examine their aggregate influence, and all relied on one-off measures of metabolic traits among middle-to-older-aged adults which gives little insight as to when in the life course metabolic alterations first occur. Examining genetic liability to type 2 diabetes in relation to repeated measures of metabolic traits starting in childhood could reveal the existence and timing of subclinical trait perturbations most central to type 2 diabetes development.

We aimed in this study to reveal early metabolic features characterising a genetic liability to adult type 2 diabetes. Using birth cohort study data, we examined genetic liability to adult type 2 diabetes in relation to detailed traits from targeted metabolomics among the same individuals at four key stages of early life – childhood (age 8y), adolescence (age 16y), young adulthood (age 18y), and adulthood (age 25y). The relative importance of traits was gauged by comparing patterns and magnitudes of genetic effects across time points. For replication analyses, two-sample Mendelian randomization [20] was also conducted in an independent sample of adults to confirm the persistence of any metabolic features of disease liability observed in early life.

## Methods

### Study populations

Data were from offspring participants of the Avon Longitudinal Study of Parents and Children (ALSPAC), a population-based birth cohort study in which 14,541 pregnant women with an expected delivery date between 1 April 1991 and 31 December 1992 were recruited from the former Avon county of southwest England [21]. Since then, 13,988 offspring alive at one year have been followed repeatedly with questionnaire- and clinic-based assessments [22], with an additional 811 children enrolled over the course of the study. Offspring were considered for the current analyses if they had no older siblings in ALSPAC (202 excluded) and of a white-European ethnicity (604 excluded) to reduce confounding of associations by high relatedness and ancestral population structure.

Written informed consent was provided and ethical approval was obtained from the ALSPAC Law and Ethics Committee and the local research ethics committee. The study website contains details of all available data through a fully searchable data dictionary and variable search tool (http://www.bristol.ac.uk/alspac/researchers/our-data/).

### Assessment of genetic liability to adult type 2 diabetes

Genotype was assessed using the Illumina HumanHap550 quad chip platform. Quality control measures included exclusion of participants with sex mismatch, minimal or excessive heterozygosity, disproportionately missing data, insufficient sample replication, cryptic relatedness, and non-European ancestry. Imputation was performed using the Haplotype Reference Consortium (HRC) panel. Since this study aims to address causation (in the direction of type 2 diabetes liability to metabolic traits), not prediction, genetic liability to type 2 diabetes was based on genetic variants associated with type 2 diabetes case status at genome-wide significance (P-value < 5.00×10^−8^) in the largest GWAS to date which identified up to 403 independent polymorphisms among adults (74,124 cases and 824,006 controls) of white-European ethnicity, explaining 18% of variance [23]. This set of variants was refined by excluding variants identified only when adjusting for body mass index and by pruning-out those variants that were in linkage disequilibrium based on R^2^ > 0.001 (retaining those SNPs with the lowest P-values) using standard packages within the MR-Base platform (http://www.mrbase.org/beta) [24]. This left 167 genetic variants highly independently associated with type 2 diabetes, 162 of which were available in imputed ALSPAC genotype data post quality control (**Supplementary (S) 1 Table**). This set was combined into a genetic risk score (GRS) using PLINK 1.9 software, specifying the effect (type 2 diabetes raising) allele and coefficient (odds ratio) from the source GWAS as external weights. Scoring was done by multiplying the number of effect alleles (or probabilities of effect alleles if imputed) at each SNP (0, 1, or 2) by its weighting, summing these, and dividing by the total number of SNPs used. The score therefore reflects the average per-SNP effect on type 2 diabetes.

### Assessment of metabolic traits

Offspring participants provided non-fasting blood samples during a clinic visit while aged approximately 8y, and fasting blood samples from clinic visits while aged approximately 16y, 18y, and 25y. Proton nuclear magnetic resonance (^1^H-NMR) spectroscopy as part of a targeted metabolomics platform [13,25] was performed using EDTA-plasma samples (stored at or below −70 degrees Celsius before processing) to quantify 229 metabolic traits (149 concentrations plus 80 ratios). These included the concentration and size of lipoprotein subclass particles and their cholesterol and triglyceride content, apolipoproteins, fatty acids, pre-glycemic factors including lactate and glucose, amino acids, ketone bodies, and inflammatory glycoprotein acetyls.

### Assessment of adiposity and type 2 diabetes status

For descriptive purposes, body mass index (BMI) was calculated at each time point as weight (kg) divided by squared height (m^2^) based on clinic measures of weight to the nearest 0.1 kg using a Tanita scale and height measured in light clothing without shoes to the nearest 0.1 cm using a Harpenden stadiometer. Type 2 diabetes status was not assessed at age 8y because blood glucose was not quantified based on fasting samples and no data were collected regarding physician diagnosis. Type 2 diabetes was defined at age 16y as a clinic fasting glucose ≥ 7 mmol/l, and at age 18y and 25y as a clinic fasting glucose ≥ 7 mmol/l or reported physician diagnosis of type 2 diabetes by that age.

### Statistical approach

In the first set of analyses, separate linear regression models with robust standard errors were used to estimate coefficients and 95% confidence intervals for associations of the type 2 diabetes GRS (in standard deviation (SD) units) with each metabolic trait measured at age 8y (also in SD units) as dependent variables, adjusted for sex and age at the time of metabolic trait assessment. These were repeated for metabolic traits measured at age 16y, 18y, and 25y. Effect estimates are interpreted within a ‘reverse Mendelian randomization’ framework [15] and are taken to reflect ‘metabolic features’ of liability to type 2 diabetes (not ‘metabolic causes’ of type 2 diabetes). To estimate the magnitude of associations at the extremes of genetic risk, models were repeated using the upper vs. lower quartile of an unstandardized type 2 diabetes GRS. For the purpose of comparing evidence of linear change in coefficients across time points, we ran separate linear mixed models utilising repeated measures of each metabolic trait and examined P-values from an interaction term between the type 2 diabetes GRS and mean age from these models at each time point. All models were additionally run using original (non-SD; mostly mmol/l) units to aid clinical interpretation and external comparisons.

To allow full use of measured data, the aforementioned analyses were first conducted using maximum numbers of participants with N varying across ages and between traits. Participants were eligible for inclusion in analyses at any age if they had data on genotype, sex, age, and at least 1 metabolic trait. This resulted in 6,218 eligible participants, including up to 4,761, 2,928, 2,612 and 2,560 participants at age 8y, 16y, 18y and 25y, respectively. Analyses were repeated using a consistent sample of participants with complete data on genotype, sex, age, and each metabolic trait at each time point. These sampling approaches are illustrated in **Figure 1**.

**Figure 1.**
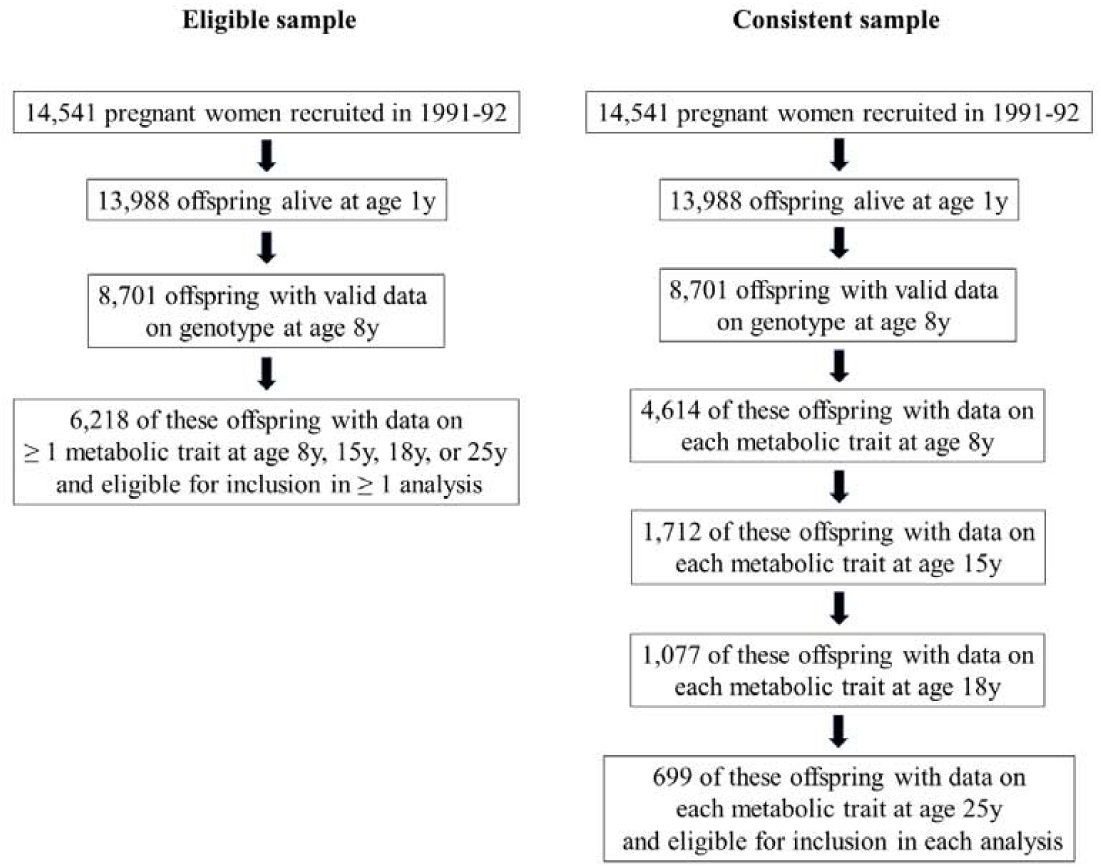
Selection of participants into analyses

### Additional analyses

We conducted two-sample Mendelian randomization (MR) analyses to estimate the metabolic features of type 2 diabetes in an independent sample of adults. The exposure in this analysis (liability to type 2 diabetes) was the 167 SNP GRS as previously described (**Supplementary (S) 1 Table**). Outcome data were GWAS summary data on 123 metabolic traits quantified using the same NMR metabolomics platform as used in present ALSPAC analyses [26], measured on between 13,476 and 24,925 adults of European ancestry [26]. Across included studies, mean (SD) age ranged from 23.9y (2.1y) to 61.3y (2.9y) and female sex ranged from 37% to 64%. Three statistical methods were used to generate MR estimates using the TwoSampleMR R package within the MR-Base platform [24]: random-effects inverse variance weighted (IVW) [24], MR-Egger [27] and weighted median [28] models, which each make differing assumptions about directional pleiotropy [29]. Consistent estimation of effect using different MR models was considered to strengthen evidence against the null and reduce the probability of type 1 errors. These estimates are interpreted as the SD-unit difference in metabolic trait per 1 log odds of type 2 diabetes.

Seventeen principal components explain 95% of the variance in these highly correlated metabolic traits based on previous ALSPAC analyses [30]. These can be used to correct nominal significance thresholds for multiple testing using the Bonferroni method (e.g. alpha=0.05/17). Since our study aims align more with (non-threshold-based) statistical estimation than (threshold-based) statistical discovery, we provide exact P-values and focus on effect size and precision [31,32]. Analyses were conducted using Stata 15.1 (StataCorp, College Station, Texas, USA).

## Results

### Sample characteristics

As shown in **Table 1**, the sample of eligible participants was 49.7% male with progressively higher BMI at later ages; mean (SD) BMI in kg/m^2^ was 16.2 (2.0), 21.4 (3.5), 22.7 (4.0), and 24.8 (4.9) at ages 8y, 16y, 18y, and 25y, respectively. Type 2 diabetes prevalence was very low across time points, with < 5 cases at age 16y, 6 cases (0.2%) at age 18y, and 7 cases (0.4%) at age 25y. Summary metabolic trait levels were generally stable across time points, with a small reduction in mean (SD) glucose from 4.2 (0.5) mmol/l at age 8y, to 3.9 (0.4) mmol/l at age 25y. Participants who were ineligible for any analysis had slightly higher BMI than those who were eligible – e.g. 25.4 kg/m^2^ vs. 24.8 kg/m^2^ at age 25y, respectively. Type 2 diabetes prevalence and summary metabolic trait levels were also comparable (**S2 Table**).

**Table 1.**
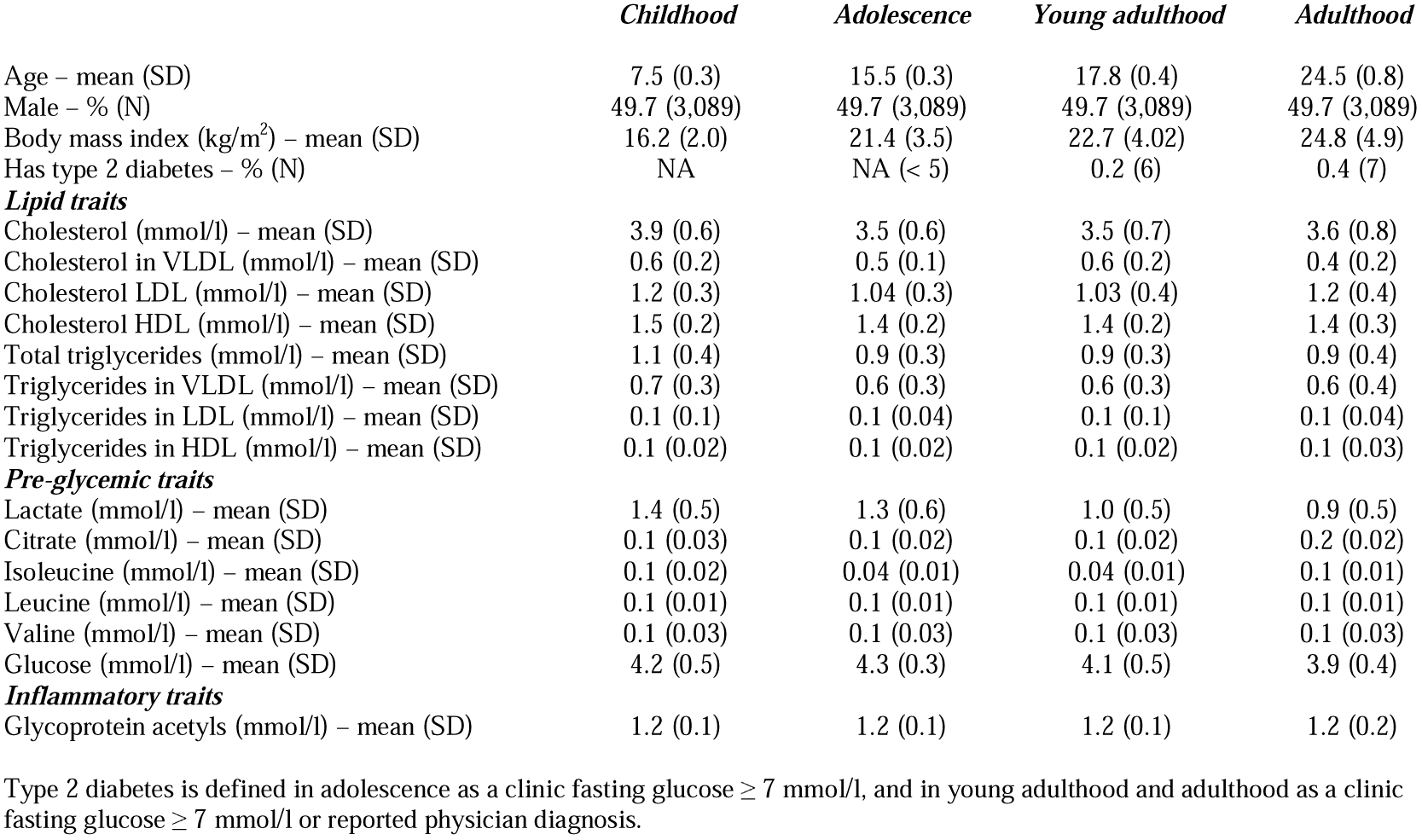
Summary metabolic traits at different early life stages among 6,218 ALSPAC offspring eligible for inclusion in ≥ 1 analysis

Of 8,701 participants with data on genotype, 4,614 additionally had data on each metabolic trait at age 8y. Of these, 1,712 additionally had data on each metabolic trait at age 16y; 1,077 of which additionally had data on each metabolic trait at age 18y. Of these, 699 participants also had data on each metabolic trait at age 25y and comprised the consistent (complete case) sample. Characteristics of this consistent sample were comparable to those of the wider sample (**S3 Table**), and differences between excluded and included participants were small **(S4 Table**).

### Associations of genetic liability to adult type 2 diabetes with metabolic traits at different early life stages in ALSPAC

At age 8y, higher type 2 diabetes liability was unassociated with lipid content in most lipoprotein particle types including very-low density lipoprotein (VLDL) with effects of inconsistent direction and magnitudes near zero. Associations were more consistent in direction and magnitude with lower cholesterol, triglycerides, and other lipids in very-large and large high-density lipoprotein (HDL) – e.g. −0.03 SD (95% CI = −0.06, −0.003; P = 0.03) for total lipids in very large HDL per SD-higher GRS (**Figure 2; S5 Table**).

**Figure 2.**
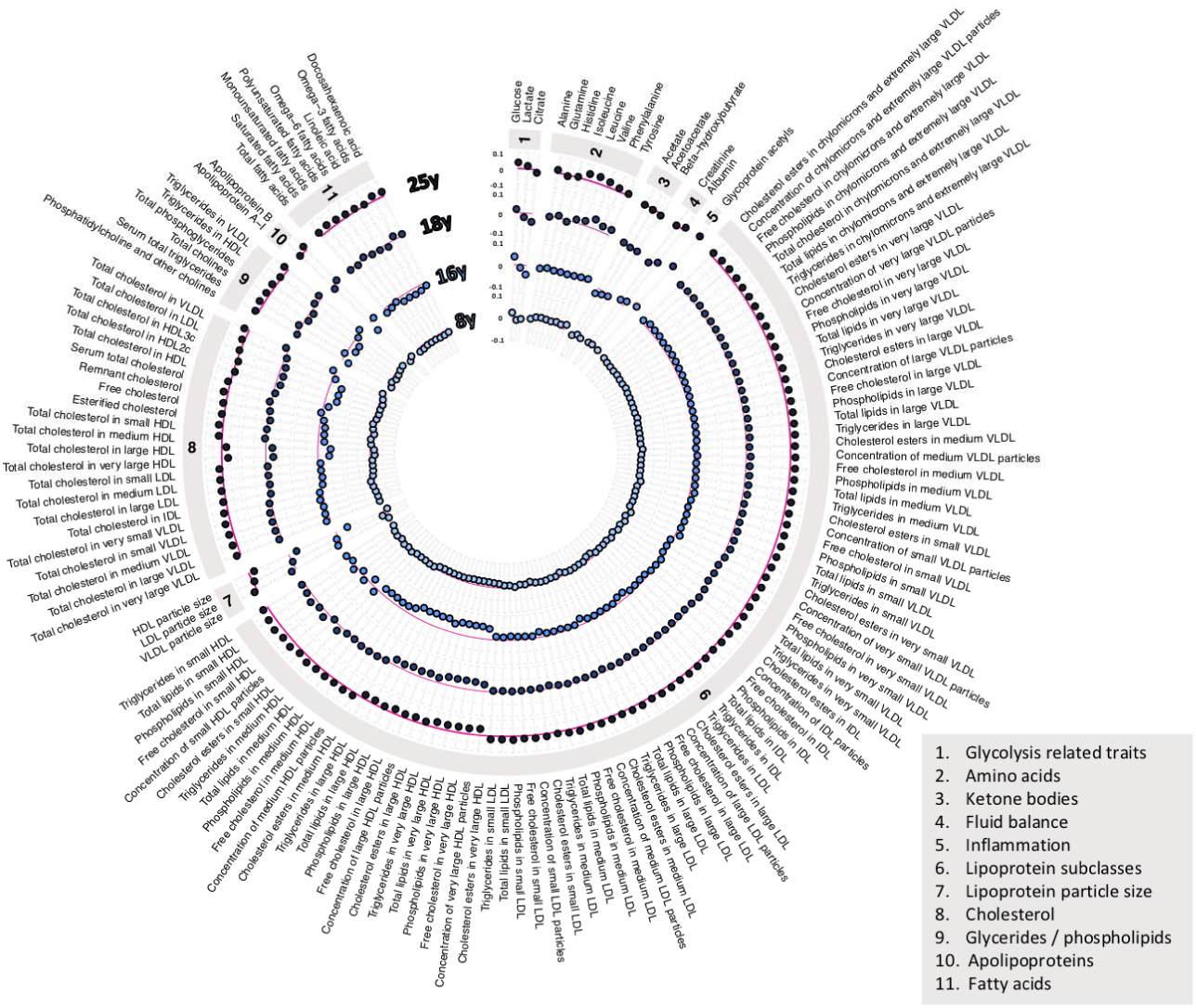
Associations of genetic liability to adult type 2 diabetes with metabolic traits at different early life stages among ALSPAC offspring. Estimates shown are beta coefficients representing SD difference in metabolic trait per SD higher GRS for type 2 diabetes, ordered concentrically (inner circle to outer circle) by increasing age at measurement.

At age 16y, higher type 2 diabetes liability (per SD-higher GRS) was weakly but more consistently associated with higher lipid content in VLDL and lower lipid content in LDL. Associations were again strongest with lower lipid content in very-large and large HDL – e.g. −0.08 SD (95% CI = −0.11, −0.04; P = 3.30×10^−05^) for total lipids in very large HDL per SD-higher GRS. Associations were also evident with pre-glycemic traits including citrate (- 0.06 SD, 95% CI = −0.09, −0.02; P = 1.41×10^−03^) and glucose (0.05 SD, 95% CI = 0.02, 0.08; P = 0.004), and glycoprotein acetyls (0.05 SD, 95% CI = 0.01, 0.08; P = 0.01).

At age 18y, higher type 2 diabetes liability (per SD-higher GRS) remained weakly associated with higher lipid content in VLDL and LDL, but more strongly associated with lower lipid content in HDL, particularly very-large and large HDL. Associations with branched chain and aromatic amino acids were also strengthened (e.g. valine: 0.06 SD; 95% CI = 0.02, 0.09; P = 1.24×10^−03^ and tyrosine: 0.04 SD; 95% CI = 0.001, 0.07; P = 0.04). Associations remained stable with glycoprotein acetyls (0.06 SD; 95% CI = 0.02, 0.10; P = 1.69×10^−03^).

At age 25y, associations had strengthened between higher type 2 diabetes liability and lipid content in VLDL subtypes such that effect size magnitudes were comparable to those seen with lipids in HDL subtypes – e.g. 0.05 SD (95% CI = 0.01, 0.09; P = 0.02) higher total cholesterol in VLDL vs. −0.06 SD (95% CI = −0.09, −0.02; P = 1.19×10^−03^) lower total cholesterol in very large HDL, each per SD-higher GRS. Increasing effect size for VLDL lipids were supported by relatively low P-values for trend across time points based on linear mixed models (**S5 Table**) – e.g. P-trend = 0.01 for total cholesterol in VLDL. These P-trend values were higher for lipids in HDL (e.g. P-trend = 0.15 for total lipids in very-large HDL), indicating more stable effect sizes across time points. Associations were also more evident with several fatty acid traits including a lower ratio of linoleic to total fatty acids (−0.07; 95% CI = −0.10, −0.03; P = 5.49×10^−04^) and lower ratios of omega-6-to-total and polyunsaturated-to-total fatty acids. Associations remained relatively strong with branched chain amino acids – e.g. with leucine at 0.06 SD; 95% CI = 0.03, 0.10; P = 8.46×10^−05^, and with glycoprotein acetyls at 0.06 SD; 95% CI = 0.01, 0.10; P = 2.96×10^−03^.

Quartile-based estimates (for the highest vs. lowest quartile of GRS) showed similar patterns with traits across time points (**S6 Table**), with effect sizes being consistently largest with lipids in very-large and large HDL – e.g. −0.08 SD at age 8y, −0.20 SD at age 16y, −0.12 SD at age 18y, and −0.13 SD at age 25y for total lipids in very-large HDL. Elevations were also consistently seen across time points for branched chain amino acids – e.g. 0.04 SD, 0.04 SD, 0.06 SD, and 0.14 SD for leucine; for glycoprotein acetyls (−0.01 SD, 0.14 SD, 0.17 SD, and 0.13 SD); and for glucose itself (0.06 SD, 0.12 SD, 0.08 SD, and 0.15 SD).

Association patterns were also comparable for standardized and quartile-based GRS estimates using a sample of 699 participants with complete data on all metabolic traits at each time point (**S7-8 Table; S1 Figure**), with effect sizes most consistently elevated for lipids in very-large and large HDL. These estimates were expectedly less precise given lower statistical power. Results based on original (non-SD) units for metabolic traits are presented in **S9-12 Table**. Mean and SD values for each metabolic trait are provided in **S13 Table** to aid effect size conversions.

### Associations of genetic liability to adult type 2 diabetes with metabolic traits in adulthood in GWAS summary data

Results of two-sample MR analyses in an independent sample of adults indicated a largely persistent pattern of associations between genetic liability to type 2 diabetes and metabolic traits seen across early life (**Figure 3; S14 Table**). Higher genetic liability to type 2 diabetes was generally positively associated with VLDL lipid subtypes and inversely associated with HDL lipid subtypes, again for large and very large HDL specifically – e.g. - SD (95% CI = −0.007, −0.002; P = 8.45×10^−04^) per 1 log odds of type 2 diabetes for total lipids in large HDL. Type 2 diabetes liability was positively associated with branched chain amino acid levels (e.g. 0.004 SD of leucine, isoleucine and valine per 1 log odds of type 2 diabetes). There was less evidence of association between type 2 diabetes liability and glycoprotein acetyls, at 0.003 SD (95% CI = 0.0001, 0.005; P = 0.04) per 1 log odds of type 2 diabetes. Glucose was the most strongly associated metabolic trait at 0.008 SD (95% CI = 0.006, 0.010; P = 5.27×10^−11^) per 1 log odds of type 2 diabetes. Evidence of effect heterogeneity was strong for most metabolic traits – e.g. Cochrane’s Q P-value=7.83×10^−16^ for the glucose IVW estimate. Where IVW estimates suggested evidence of a causal association, weighted median estimators were consistent whereas MR Egger estimates were imprecise, although there was little evidence to suggest that MR Egger intercept estimates differed from zero for any of the metabolic traits (P > 0.003).

**Figure 3.**
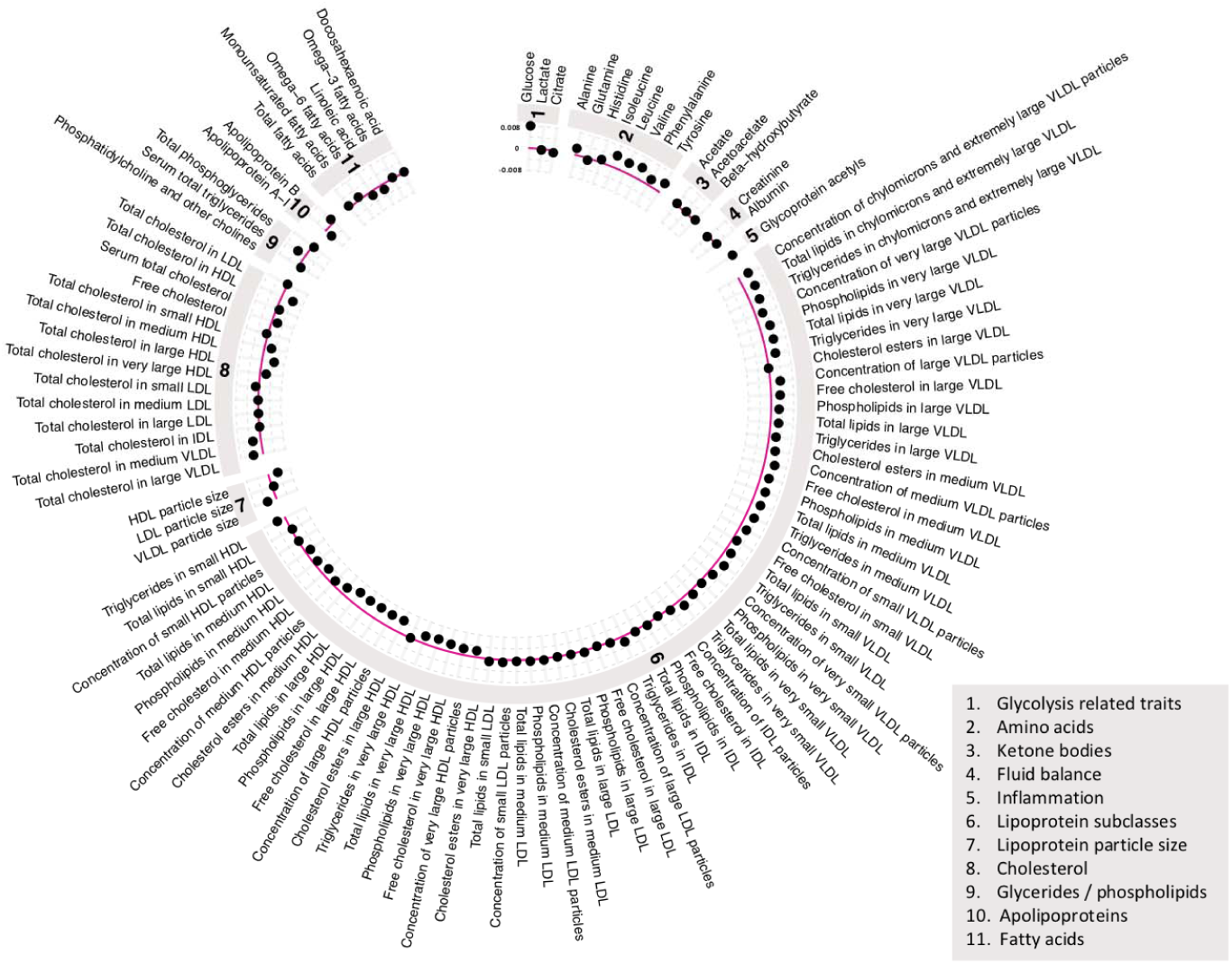
Associations of genetic liability to adult type 2 diabetes with metabolic traits in an independent sample of adults based on two-sample MR. Estimates shown are beta coefficients representing the SD-unit difference in metabolic trait per 1 log odds of type 2 diabetes based on the IVW method.

## Discussion

This study aimed to reveal early metabolic features of type 2 diabetes liability by integrating genetic liability to adult type 2 diabetes with detailed metabolic traits measured across early life (from age 8y to 25y). These metabolic traits were measured long before the expected clinical onset of type 2 diabetes [33], and consequently, their perturbations are expected to reflect early signs of disease that are detectable in circulation. Our findings suggest that one of these earliest features is lower lipid content in HDL – particularly in large and very large particle subtypes – which precedes lower citrate and higher branched chain amino acid and inflammatory glycoprotein acetyl levels. This feature is apparent in childhood as early as age 8y, several decades before the clinical onset of disease. Persistent patterns of effect were observed in an independent sample of adults based on two-sample MR, supporting their continued relevance with advancing age.

Adiposity is expected to be a key driver of type 2 diabetes and its metabolic intermediates, with several MR [20] studies supporting profound effects of higher BMI on metabolic dysfunction [34,35] and clinical type 2 diabetes in adulthood [4,35]. More specifically, higher BMI is strongly associated with lower cholesterol in HDL and higher cholesterol in more atherogenic lipid types including VLDL, remnant particles, and apolipoprotein-B; higher branched chain amino acid levels, and higher glycoprotein acetyls – a marker of chronic inflammation [36] heavily implicated in type 2 diabetes pathogenesis via impaired secretion and sensitivity of insulin [37,38]. These effects are evident in early adulthood (mean age 24y) [34] and likely also in childhood, with recent observational findings in ALSPAC suggesting these same patterns of effect at age 10y [39]. Visceral fat mass is in turn expected to drive metabolic effects of total adiposity, with recent observational findings based on NMR metabolomics in the Netherlands Epidemiology of Obesity (NEO) study suggesting that lipids in HDL are inversely associated, while lipids in VLDL are positively associated, with visceral fat volume [40]. As seen presently in response to type 2 diabetes liability, HDL associations were strongest for very-large and large particle subtypes, while branched chain amino acid and glycoprotein acetyl levels also showed similar patterns of association in NEO in relation to visceral fat volume as seen here in response to type 2 diabetes liability, supporting visceral fat as a common influencer.

Pressing questions concern which of these BMI-induced intermediates are causes of type 2 diabetes, and which arise in response to the developing disease process – i.e. are part of type 2 diabetes itself. One recent MR study suggested that higher branched chain amino acid levels are associated with higher type 2 diabetes risk, implying that perturbed branched chain amino acid metabolism causes type 2 diabetes [41]. However, instrumenting metabolic traits is difficult in an MR framework because their genetic architecture overlaps greatly [42,43], resulting in use of common genetic variants discovered in GWAS that are not typically specific to one metabolic trait. Using the same genetic variant for multiple traits outside of a multivariable MR framework [44] can lead to inflated MR estimates via a ‘double-counting’ of allele effects, as was likely the case in that study [41,43] where the same single genetic variant (rs1440581) was used as an instrument for both leucine and valine.

In contrast, a more recent MR study suggested that higher genetic liability to insulin resistance – a precursor of type 2 diabetes – raises branched chain amino acid levels [45], positioning these as consequences of type 2 diabetes development, a conclusion further supported by present results based on a GRS for type 2 diabetes itself and metabolomics data across early and adult life. Elevated branched chain amino acid levels are attributed to an inability to properly suppress protein breakdown in the insulin resistant state [46], and results of experimental studies in humans suggest that infusions of branched chain amino acids acutely worsen insulin sensitivity [47-50]. Experimental animal studies suggest that in the context of high fat consumption, increased dietary intake of branched chain amino acids contributes to the development of obesity-associated insulin resistance [51], while pharmacological activation of branched chain amino acid dehydrogenase complex – the second step in branched chain amino acid catabolism – leads to increased branched chain amino acid disposal and improved insulin sensitivity in Zucker rats [52].

In that same MR study [45], strong effects of higher insulin resistance were also found on higher total-lipid content in VLDL subtypes; on lower total lipid content in very-large, large, and medium HDL subtypes; on higher total triglycerides and apolipoprotein-B; and on lower beta-hydroxybutyrate and citrate – patterns much like those seen presently. Effect sizes were, however, much larger in previous results. This is likely because insulin resistance is more biologically distinct than type 2 diabetes (a highly heterogenous disease) and likely has more precise metabolic effects. Differences in effect size also likely reflect the much younger ages at which traits were measured in the present study; effects of the disease process are expected to strengthen over time. Effect sizes pertaining to the earliest stages of disease are expected to be small, but their patterns and relative magnitudes are nevertheless informative. For example, perturbations were detected early for circulating citrate – an important substrate of cellular energy metabolism that has been shown to dramatically increase in people with rapid diabetes improvement following bariatric surgery [53]. Citrate synthase activity is also impaired in myotubes from people with type 2 diabetes [54].

Bi-directional effects between metabolic traits and liability to type 2 diabetes remain plausible, but larger GWAS and methodological advancements are needed for handling overlapping genetic instruments to confirm this. Additional clues come from another recent MR study that examined effects of type 2 diabetes liability on metabolic traits derived from NMR and mass-spectrometry platforms; and effects in the reverse direction for 20 instrumentable traits [19]. Genetically higher cholesterol in HDL – again in very-large and large subtypes – were most strongly associated with lower fasting glucose, but these effects did not extend to type 2 diabetes itself. Metabolic traits with the strongest genetic evidence of effect on type 2 diabetes were phospholipids in VLDL and intermediate-density lipoprotein (IDL) particles, and total triglycerides [19]. Conversely, type 2 diabetes genetic liability had the greatest effect on the amino acid alanine, along with several phosphatidylcholine alkylacyls, providing further evidence that such perturbations are more a consequence than a cause of type 2 diabetes liability.

Importantly, results of both recent MR studies [19,45] were based solely on samples of middle-to-late aged adults, which are useful for examining clinical disease endpoints and their intermediates, but are on their own less useful for examining the timing of their onset. Our results based on younger participants affirm the patterns seen among older adults [19,45] and shed light on the temporal nature of type 2 diabetes development, suggesting that perturbed HDL cholesterol metabolism is one of the first events in the developmental stages of type 2 diabetes (a consequence of processes leading to the disease), even before perturbations in VLDL cholesterol, triglycerides, branched chain amino acids, or inflammation become apparent. The patterns of effect seen amongst young people here were also seen in an independent sample of adults based on two-sample MR, with higher genetic liability to type 2 diabetes related to lower lipids in large and very large HDL subtypes, higher lipids in VLDL, and higher branched chain amino acid levels. The particularly early tracking of HDL lipids suggests that HDL metabolism aligns most closely with adipose tissue insulin resistance and may be a reliable early marker of impending type 2 diabetes.

### Limitations

Limitations of this study include modest sample sizes for ALSPAC analyses, particularly for consistent (complete case) analyses. This stems from the rarity of participation at four clinic occasions spanning childhood to adulthood and the need for data on an extensive set of variables. Descriptive comparisons were made for key measured traits between excluded and included participants, and these differences appeared small – e.g. BMI was 25.0 kg/m^2^ vs. 24.5 kg/m^2^ at age 25y in the compete-case sample, respectively. The metabolic traits examined had limited coverage of pre-glycemic, protein, and hormonal traits; insulin was notably absent as it is not quantified by NMR and was not available at the first time point. Six metabolic traits (diacylglycerol, ratio of diacylglycerol to triglycerides, fatty acid chain length, degree of unsaturation, conjugated linoleic acid, and ratio of conjugated linoleic acid to total fatty acids) were not measured at the final age 25y time point; however these traits were analysed at earlier time points to meet the aim of comparing early perturbances. Blood samples from the first occasion of metabolic trait assessment were derived while non-fasting, but trait concentrations have shown stability over different durations of fasting time in previous analyses [55]. Diagnoses of type 2 diabetes in ALSPAC were based on fasting plasma glucose or reported physician diagnosis; glycated haemoglobin (HbA1c) measures were not available. However, recommendations by the American Diabetes Association in 2018 note the equal appropriateness of fasting plasma glucose and HbA1c as diagnostic traits for type 2 diabetes [6], and prevalence is expected to be low at the young ages studied regardless of the criteria used.

Our analyses were restricted to white-Europeans which helps to reduce confounding by ancestral population structure, but limits inference to other groups. This requires more comprehensive GWAS studies of non-white-European populations together with metabolomic measurements in cohort studies with higher representation of those groups. Extending analyses to other populations with high present and future burdens of type 2 diabetes, such as African and east-Asian populations [1], is vital for understanding the early features of type 2 diabetes in these groups and how disease burden could be mitigated. Although our two-sample MR analysis confirmed the same metabolic features in an independent sample, differences in the units of exposure (measured as ‘per SD higher GRS for type 2 diabetes’ in ALSPAC vs. ‘per 1 log odds of type 2 diabetes’ in summary GWAS data) prevent direct comparison of effect sizes between these analyses; however patterns of effect across metabolic traits are still considered informative. We aimed to reveal early metabolic features of type 2 diabetes, but objectives are only strictly feasible within a framework of liability to type 2 diabetes because the current study population is without clinical disease. This was deliberate on our part as a novel approach for identifying early features of disease based on genetic susceptibility to adult disease and repeat metabolomics in early life, with implications for pre-clinical populations. Lastly, although we primarily interpret results as reflecting the metabolic effects of type 2 diabetes liability, alternative scenarios including bias or pleiotropy remain possible explanations (7 such scenarios are proposed by Holmes and Davey Smith [15]). Directly interrogating each scenario remains difficult, but methodological flexibility together with an increasingly large scale and scope of genomic and metabolomic data should make doing so more feasible in the near future.

### Conclusions

Our results based on genetic liability to adult type 2 diabetes in relation to repeated measures of detailed metabolic traits across early life suggest that one of the earliest metabolic features of type 2 diabetes liability is lower lipid content in HDL particles – particularly in very-large and large HDL subtypes – which precedes other perturbations including lower citrate and higher branched chain amino acid and inflammatory glycoprotein acetyl levels. This feature is apparent in childhood as early as age 8y, several decades before the clinical onset of disease.

## Conflicts of interest

None to declare.

## Acknowledgments

We are grateful to the families who participated in this study, the midwives for their help in recruiting them, and the entire ALSPAC team which includes interviewers, computer and laboratory technicians, clerical workers, research scientists, volunteers, managers, receptionists, and nurses.

## Funding

The UK Medical Research Council, Wellcome (102215/2/13/2), and the University of Bristol provide core support for ALSPAC. GWAS data was generated by Sample Logistics and Genotyping Facilities at Wellcome Sanger Institute and LabCorp (Laboratory Corporation of America) using support from 23andMe. This publication is the work of the authors who are guarantors for its contents. JAB is supported by Cancer Research UK (C18281/A19169) and the Elizabeth Blackwell Institute for Health Research, University of Bristol and the Wellcome Trust Institutional Strategic Support Fund (204813/Z/16/Z). EEV and CB are supported by Diabetes UK (17/0005587). NJT is a Wellcome Trust Investigator (202802/Z/16/Z), is the PI of the Avon Longitudinal Study of Parents and Children (MRC & WT 102215/2/13/2), is supported by the University of Bristol NIHR Biomedical Research Centre (BRC-1215-20011), the MRC Integrative Epidemiology Unit (MC_UU_12013/3) and works within the CRUK Integrative Cancer Epidemiology Programme (C18281/A19169). DC and GDS work in a unit funded by the UK Medical Research Council (MC_UU_00011/1,6) and the University of Bristol. The funders had no role in study design, data collection and analysis, decision to publish, or preparation of the manuscript.

## Author contributions

All authors contributed to the planning of the study. JAB and CJB had access to data and conducted analyses. JAB, CJB, and EEV wrote the first draft. All authors critically reviewed the intellectual content of manuscript drafts and approved the final version for submission.

---

**S1 Figure.**
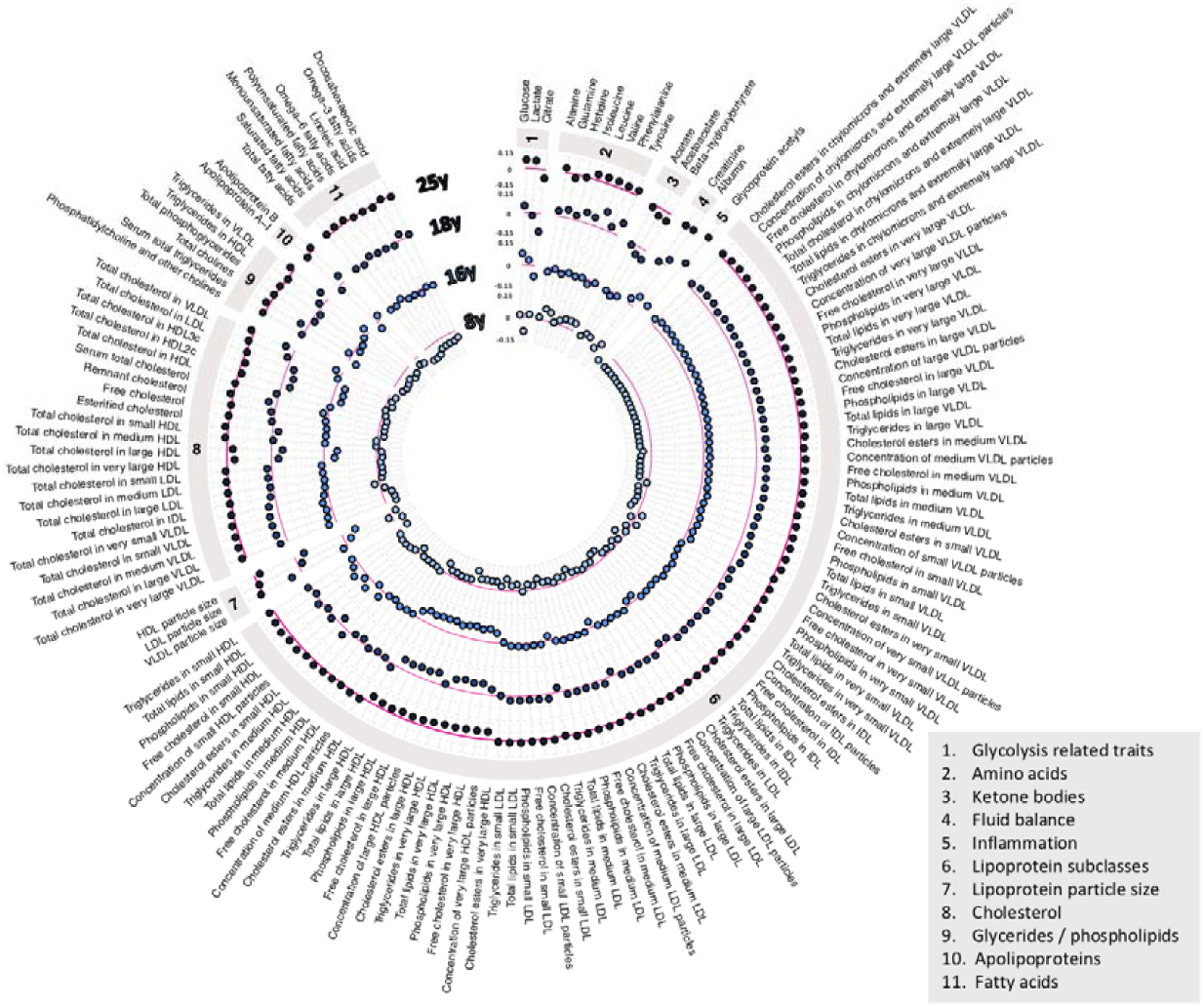
Associations of genetic liability to adult type 2 diabetes with metabolic traits at different early life stages among a consistent sample of 699 ALSPAC offspring. Estimates shown are beta coefficients representing SD difference in metabolic trait per SD higher GRS for type 2 diabetes, ordered concentrically (inner circle to outer circle) by increasing age at measurement.

